# Single-domain antibodies reveal unique borreliacidal epitopes on the Lyme disease vaccine antigen, Outer surface protein A (OspA)

**DOI:** 10.1101/2024.01.23.576890

**Authors:** David J Vance, Saiful Basir, Carol Lyn Piazza, Graham Willsey, H M Emranul Haque, Jacque M Tremblay, Michael J Rudolph, Beatrice Muriuki, Lisa A Cavacini, David D Weis, Charles B Shoemaker, Nicholas J Mantis

## Abstract

Camelid-derived, single-domain antibodies (V_H_Hs) have proven to be extremely powerful tools in defining the antigenic landscape of immunologically heterogeneous surface proteins. In this report, we generated a phage-displayed V_H_H library directed against the candidate Lyme disease vaccine antigen, Outer surface protein A (OspA). Two alpacas were immunized with recombinant OspA serotype 1 (ST1) from *Borrelia burgdorferi* sensu stricto strain B31, in combination with the canine vaccine RECOMBITEK^®^ Lyme containing lipidated OspA. The phage library was subjected to two rounds of affinity enrichment (“panning”) against recombinant OspA, yielding 21 unique V_H_Hs within two epitope bins, as determined through competition ELISAs with a panel of OspA-specific human monoclonal antibodies. Epitope refinement was conducted by hydrogen exchange-mass spectrometry (HX-MS). Six of the monovalent V_H_Hs were expressed as human IgG1-Fc fusion proteins and shown to have functional properties associated with protective human monoclonal antibodies, including *B. burgdorferi* agglutination, outer membrane damage, and complement-dependent borreliacidal activity. The V_H_Hs displayed unique reactivity profiles with the seven OspA serotypes associated with *B. burgdorferi* genospecies in the United States and Europe consistent with there being conserved epitopes across OspA serotypes that should be considered when designing and evaluating multivalent Lyme disease vaccines.

## INTRODUCTION

Lyme disease is the most common vector-borne infection in the United States, with an estimated 450,000 cases per year (1). The primary etiologic agent of Lyme disease in the US is the spirochetal bacterium, *Borrelia burgdorferi* sensu stricto (herein referred to as simply *B. burgdorferi*) with other genospecies responsible for disease in Europe and Asia. The spirochete is transmitted to humans by the black legged tick, *Ixodes scapularis* in the eastern US and *Ixodes pacificus* in the western part of the country. During the course a blood meal, *B. burgdorferi* migrates from the tick midgut, where it normally resides, to the salivary glands where it is deposited into the skin of a host. In humans, the spirochete proliferates at the site of the tick bite, typically resulting in an expanding skin lesion commonly referred to as a bull’s eye rash or erythema migrans (2, 3). In the absence of antibiotic intervention, *B. burgdorferi* disseminates to peripheral tissues, organs, large joints, and central nervous system, potentially resulting in severe complications including neuroborreliosis, carditis and/or Lyme arthritis weeks, months or even years later (2, 4).

A myriad of Lyme disease vaccine candidates in various stages of preclinical and clinical development are focused on a single *B. burgdorferi* antigen known as outer surface protein A (**OspA**) (5–14). OspA is a ∼31 kDa lipoprotein expressed by *B. burgdorferi* during habitation of the tick midgut, then down regulated during transmission or upon entry into a vertebrate host (15). Decades ago, it was recognized that OspA vaccination or passive transfer of OspA antisera prevented *B. burgdorferi* infection in mouse and guinea pig models of tick-mediated transmission (16–20). Subsequent studies demonstrated that OspA antibodies inhibit one or more steps in spirochete migration from tick-to-mammal, although the exact mechanism(s) by which antibodies interrupt this process have not been fully elucidated (21–23). Nonetheless, recombinant OspA vaccines proved highly efficacious in Phase III clinical trials and one (LYMErix™) was licensed in the United States from 1998 to 2002 before being discontinued (14, 24, 25).

Despite ongoing investments in next-generation OspA vaccines, there remains a considerable gap in our understanding of the regions (epitopes) on OspA responsible for eliciting protective immunity to *B. burgdorferi* (26, 27). This was not necessarily an impediment with the first generation OspA vaccines like LYMErix™ because they were based on the single OspA serotype (ST1) that predominates in the United States. Globally, however, there are at least seven OspA serotypes associated with Lyme disease-causing *B. burgdorferi* genospecies: *B. burgdorferi* (ST 1), *B. afzelii* (ST 2), *B. garinii* (ST 3, 5, 6, 7) and *B. bavariensis* (ST 4) (28, 29). Concerns about the lack of serotype cross protection has prompted the engineering of hexavalent and heptavalent vaccines consisting of recombinant full length and truncated OspA derivatives (10, 11, 30, 31). An alternative strategy would be to identify common or even conserved epitopes across the two or more of the OspA serotypes and use this information in the rational design of novel vaccine antigens (32). With this goal in mind, we recently generated an epitope map of OspA ST 1 using a collection of borreliacidal and transmission blocking human and mouse monoclonal antibodies (MAbs) revealing four distinct epitope bins or clusters (32, 33). Structural analysis of a subset of MAbs in complex with OspA ST1 has revealed detailed information about the nature of a select number of protective epitopes within these bins (26, 27, 34, 35).

Camelid-derived, single-domain antibodies, technically known as V_H_Hs, have emerged as tools to define the antigenic landscape of immunologically heterogeneous surface proteins, including notoriously polymorphic influenza virus hemagglutinin and the SARS-CoV-2 RBD (36–38). V_H_Hs derive from heavy chain-only antibodies (HCAbs) that exist with the Camelidae family, including llamas and alpacas (39). HCAbs consist of two heavy chains (homodimers) without light chain partners. The terminal V_H_ domain or V_H_H confer antigen binding activity with properties and affinities not dissimilar to conventional IgG (40). V_H_Hs are small, stable and amenable to expression on the surface of bacteriophage M13 (“phage display”), thereby allowing antibody panning and affinity enrichment against targets of interest. In this report, we generated a phage-displayed V_H_H library directed against OspA ST1 and identified 21 unique V_H_Hs within two epitope bins. Six of the monovalent V_H_Hs were expressed as human IgG1-Fc fusion proteins and been shown to have functional properties associated with protective human monoclonal antibodies, including *B. burgdorferi* agglutination, outer membrane damage, and complement-dependent borreliacidal activity. Finally, the V_H_Hs displayed unique reactivity profiles with the seven OspA serotypes associated with *B. burgdorferi* genospecies in the United States and Europe consistent with there being conserved epitopes across OspA serotypes that should be considered when designing and evaluating multivalent Lyme disease vaccines.

## RESULTS

### Isolation of distinct families of OspA-specific V_H_Hs

A phage-displayed V_H_H library was constructed from two alpacas that had been immunized with recombinant OspA (rOspA) serotype 1 (ST1) from *B. burgdorferi* strain B31 in combination with the canine vaccine RECOMBITEK^®^ Lyme, which contains lipidated OspA. The resulting phage library was subjected to 2 rounds of panning against rOspA, as described in the Materials and Methods. In total, 74 phages were isolated and subjected to DNA sequencing, revealing 21 unique clones (**Table 1**). Thirteen of the 21 clones were assigned to five different families based on complementarity determining region 3 (CDR3) sequence similarities. As an example, the alignment of the four members of the L8H8 family, along with the inferred germline V_H_ and J_H_ genes, is shown in **Figure S1**. The remaining 8 V_H_Hs were designated as “orphans” because they had no significant amino acid similarity to other V_H_Hs. Sequence analysis further suggests that 20 of 21 V_H_Hs likely derived from V_H_ 3-3 germline, with the exception (L8H7) derived from V_H_ 41 (41). From a structural standpoint, V_H_ 3-3-derived antibodies tend to have CDR3 elements that are pinned back towards the core of the antibody (42). The V_H_Hs were cloned into a pET32b-based vector and expressed as E-tagged thioredoxin fusion proteins in *E. coli* Rosetta Gami 2 cells. All 21 V_H_Hs bound to rOspA by ELISA with EC_50_ values that ranged from 1.8 nM to >1 µM (**Table 1**; **Figure 1**). V_H_H dissociation constants (K_D_) ranged from 0.28 to 457 nM, as determined by biolayer interferometry (**Table 1**; **Figure S2-S3**).

**Figure 1.**
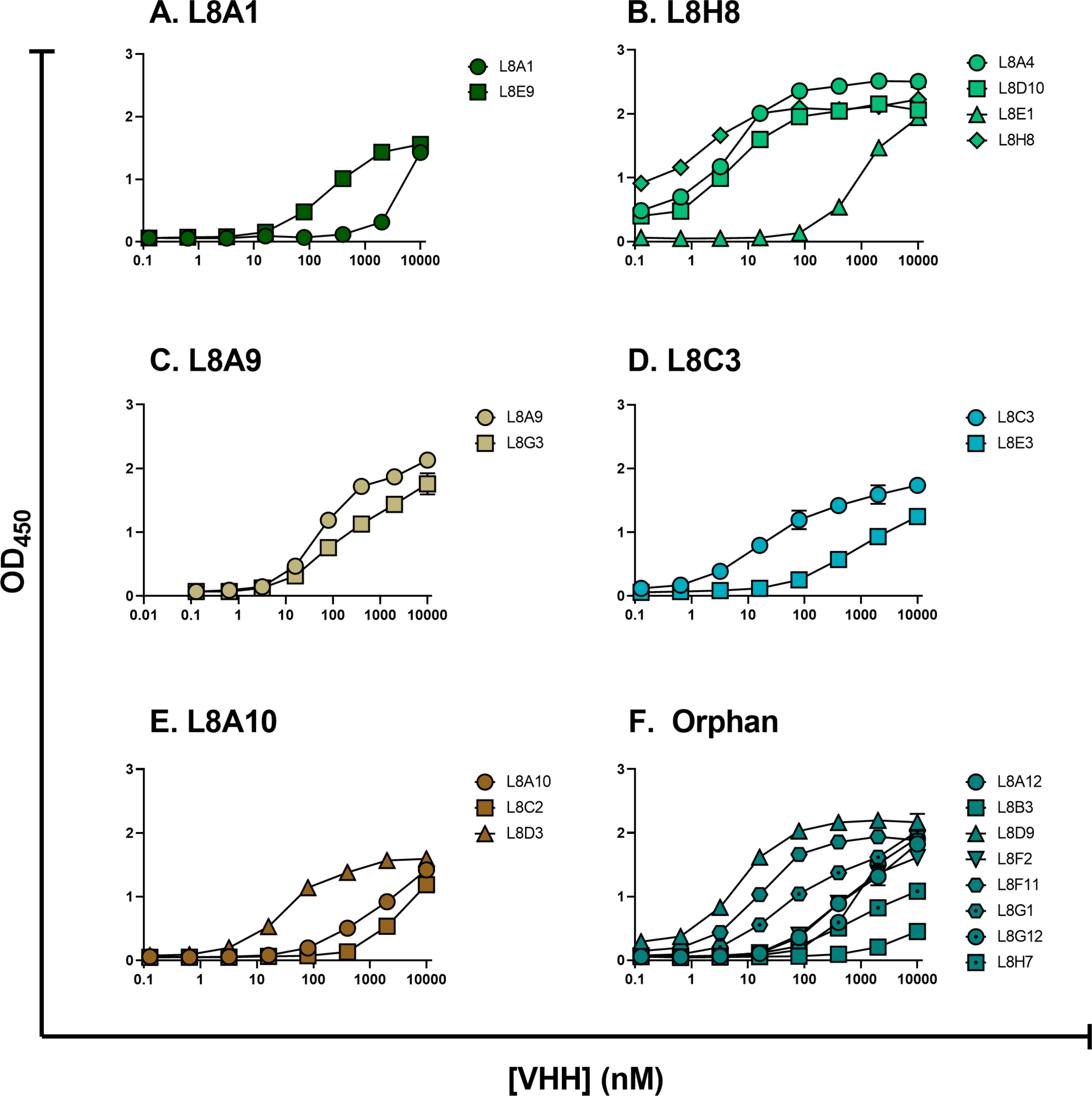
V_H_H recognition of recombinant OspA. Expressed and purified V_H_Hs were tested for binding to immobilized recombinant OspA ST1 by ELISA. V_H_Hs were grouped by clonal relationships. (A) L8A1 family; (B) L8H8 family; (C) L8A9 family; (D) L8C3 family; (E) L8A10 family; (F) “orphan” V_H_Hs not part of a clonal family.

**Table 1.**
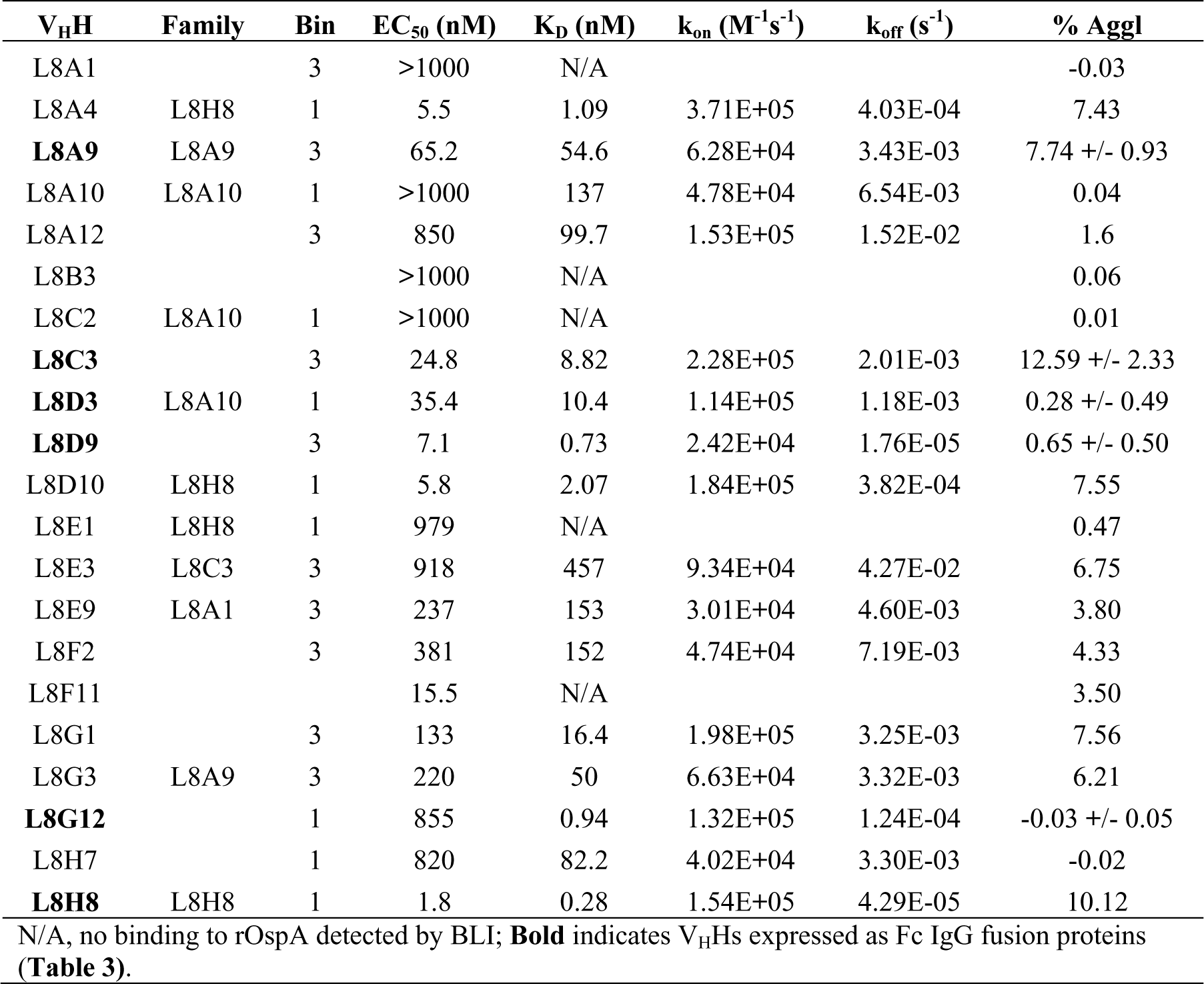
Anti-OspA V_H_Hs, families, epitope bins, binding affinities and agglutination activities.

### V_H_Hs cluster within two epitope bins on OspA

Four spatially distinct epitope bins (0-3) have been described along the length of OspA ST1 (33). Bin 0 constitutes a region of the N-terminus of OspA that is normally buried in the bacterial outer membrane and only accessible when OspA is released from the cell surface (26, 43). Bins 1 and 2 encompass OspA’s central β-sheet (strands 8-13), while Bin 3 is situated within OspA’s C-terminus (β-strands 16-21) and projects outward from the bacterial surface. MAbs capable of blocking *B. burgdorferi* tick to mouse transmission have been identified in bins 1, 2 and 3 (32, 33).

To epitope map the new panel of antibodies, V_H_Hs representative of the different clonal families were subjected to competitive BLI with MAbs from bin 1 (857-2), bin 2 (212-55), and bin 3 (LA-2) (**Figure S4**). Of the 11 representative V_H_Hs tested, four (L8H8, L8D3, L8H7, L8G12) were assigned to bin 1, based on competition with 857-2, and seven were assigned to bin 3, based on competition with LA-2 (**Table 1**; **Figure 2**). The seven V_H_Hs in bin 3 were subjected to further competition with MAbs 3-24 and 319-44, which recognize epitopes flanking LA-2. Specifically, LA-2 targets the three exposed loops between β-strands 16-17, 18-19, and 20-21, as well as part of the C-terminal α-helix [**PDB ID** 1FJ1]. 3-24 targets β-strands 16-18, while 319-44 recognizes β-strands 19, 20, and 21, along with the loops between β-strands 16-17, 18-19, and 20-21 [**PDB ID** 7T25]. The seven V_H_Hs competed with 3-24, but not 319-44, thereby positioning their epitopes in proximity to β-strands 16-18 (pink shading; **Figure 2B**).

**Figure 2.**
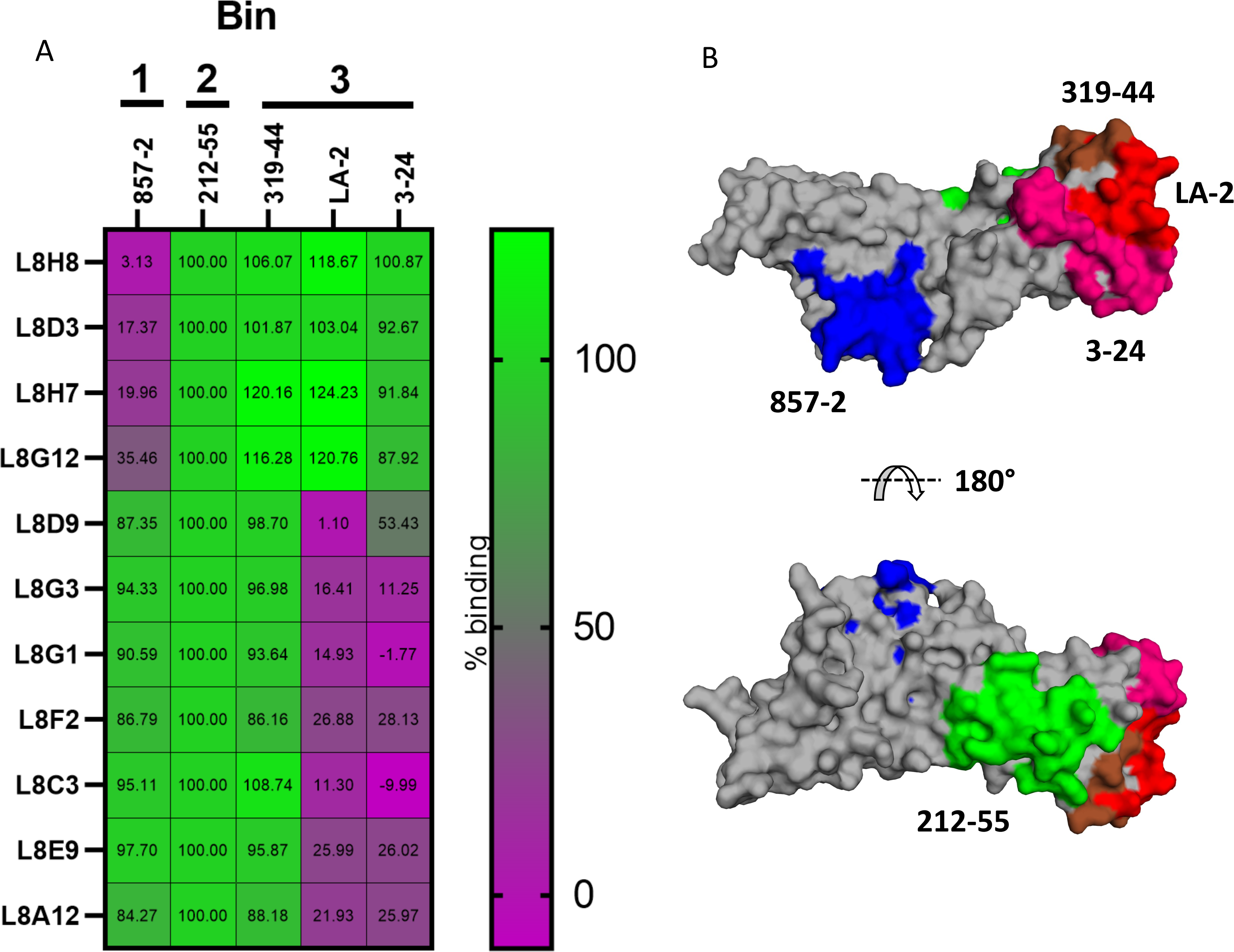
Assignment of V_H_Hs to OspA epitope bins. Representative V_H_H family members (and all orphans) were tested for their ability to bind to rOspA that had been previously coated by Bin 1-3 mAbs using BLI. A) Biotinylated-OspA was immobilized on Streptavidin coated biosensors and then coated with mAbs from Bins 1-3 (top). Individual V_H_Hs were then allowed to bind, and the change in the signal induced by the V_H_Hs is reported in each box. The data is normalized to each V_H_Hs binding in the presence of bin 2 mAb 212-55, which did not block any of the V_H_Hs from binding, and colorized so that dark purple represents strong competition/inhibition and light green represents no competition. B) Surface representation of OspA (PDB ID: 1OSP) (gray) colored with the HX-MS determined epitopes of the mAbs used in panel A) colored blue, green, and shades of red for bins 1, 2 and 3, respectively.

To better resolve antibody-OspA interactions within Bin 1, five V_H_Hs were subjected to epitope mapping by hydrogen exchange (HX)-mass spectrometry (MS) analysis using protocols optimized for OspA (33, 44). HX-MS is an increasingly powerful tool for epitope mapping in which the HX reaction is conducted in solution and captures antibody-induced changes in antigen backbone flexibility; strong reductions in HX are interpreted as points of antigen-antibody contact (45). As predicted, HX-MS indicated that all five V_H_Hs in bin 1 protected regions within OspA’s central β-sheet (33). Specifically, L8H8 and two of its family members, L8A4 and L8D10, protected nearly identical OspA peptides corresponding to β-strands 10-14 (**Table 2**; **Figure 3**). The “orphan” V_H_H, L8G12, also protected β-strands 11-14, while L8D3 (a member of the L8A10 family) protected β-strands 11-13 (**Table 2**; **Figure 3**). These results demonstrate that β-strands 10-14 constitute a V_H_H “hotspot” within bin 1.

**Figure 3.**
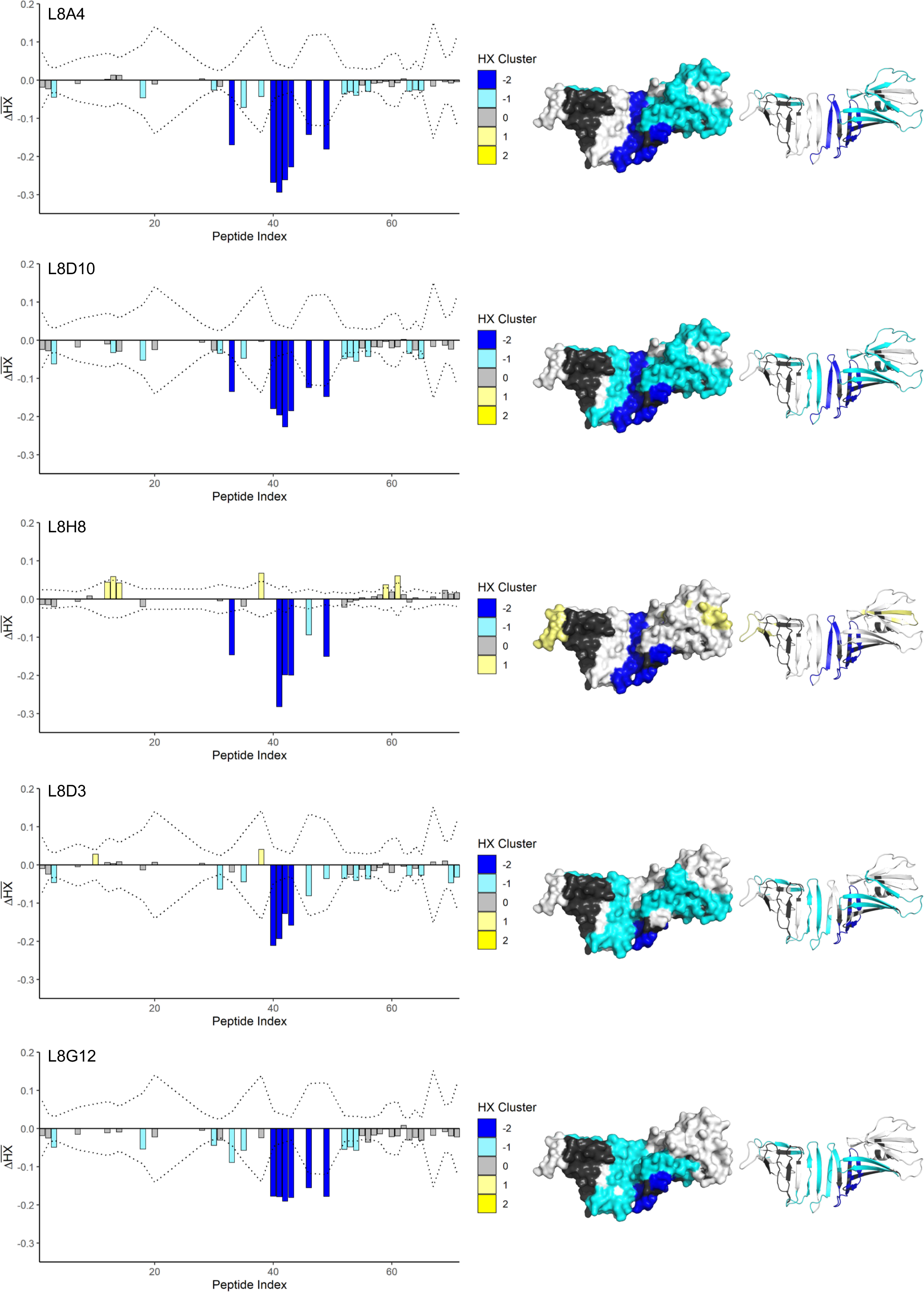
Mapping of V_H_H epitopes of OspA by HX-MS. The left hand column shows Δ*HX̅* for each individual OspA peptide, arranged from N- to C-terminals of OspA. The dotted line denotes the limit for statistically significant differences. The results were classified into categories of protection, denoted by blue and cyan for protection (slower hydrogen exchange) and yellow (faster hydrogen exchange) for the bound form of OspA relative to the free form. The center and right-hand panels show the results mapped onto the surface and ribbon representations of OspA (PDB ID: 1OSP).

**Table 2.** OspA V_H_H epitope mapping by HX-MS.

| **Table 2. OspA V<sub>H</sub>H epitope mapping by HX-MS** |  |  |  |  |
| --- | --- | --- | --- | --- |
| **V<sub>H</sub>H** | **peptides<sup>a</sup>** | **2° struct.** | **Molar ratio<sup>b</sup>** | **HX time-course<sup>c</sup>** |
| L8A4<sup>d</sup> | 129-145, 148-157, 159,<br>161-171, 174, 175, 177 | β-strands 10-14 | 4:1 | screening |
| L8D10<sup>d</sup> | 129-145, 148-157, 159,<br>161-171, 174, 175, 177 | β-strands 10-14 | 4:1 | screening |
| L8H8<sup>d</sup> | 129-145, 148-157, 159,<br>161-171, 174, 175, 177 | β-strands 10-14 | 4:1 | complete |
| L8D3 | 148-157, 159, 161-171 | β-strands 11-13 | 8:1 | screening |
| L8G12 | 148-157, 159, 161-171,<br>174, 175, 177 | β-strands 11-14 | 4:1 | screening |
| <sup>a</sup>, strongly protected peptides listed as OspA residue numbers (see supplementary information); <sup>b</sup>, molar ratio V<sub>H</sub>H:OspA; <sup>c</sup>, as described in (44); <sup>d</sup>, derive from the same clonal family. |  |  |  |  |

### Functional activity associated with OspA-specific monovalent V_H_Hs and bivalent V_H_H-IgG constructs

A hallmark of OspA antibodies, and certain monovalent Fabs, is their capacity to induce agglutination of *B. burgdorferi* in culture (46–48). We have postulated that agglutination explains, at least in part, how OspA antibodies entrap *B. burgdorferi* within the tick midgut and block transmission to vertebrate hosts (46). We employed a quantitative flow cytometry-based assay to examine whether any of the 21 monovalent OspA V_H_Hs induced agglutination of live *B. burgdorferi* B31 (46). The handful of V_H_Hs with dissociation constants ≥1000 nM did not promote spirochete agglutination (**Table 1**). The remaining V_H_Hs displayed a range of agglutinating activities, although with no clear relationship between agglutination and epitope specificity (i.e., bin 1 or bin 3) or agglutination and OspA binding affinity (K_D_). For example, L8D9 (bin 3) and L8G12 (bin 1) each have sub-nanomolar binding affinities for OspA, but neither induced notable agglutination of *B. burgdorferi* (**Table 1**). L8D3 (bin 1) and L8C3 (bin 3) have comparable affinities for OspA (10.4 vs 8.82 nM), but L8D3 is a poor agglutinator while L8C3 was a good agglutinator (0.28% vs 12.5%).

To investigate the impact of antibody avidity on spirochete agglutination, three V_H_Hs each from bin 1 and bin 3 were grafted onto human IgG1 Fc elements and expressed as bivalent molecules in Expi293 cells (49). The six V_H_H-IgG fusion proteins recognized native OspA on the surface of live *B. burgdorferi* B31 to levels similar to LA-2, as demonstrated by flow cytometry (**Table 3**; **Figure 4**). The V_H_H-IgGs also had *B. burgdorferi* agglutinating activities that were significantly greater than their monovalent V_H_H counterparts (**Table 4**). For example, V_H_H L8D3 (bin 1) had <1% agglutinating activity as a monomer but >17% activity as a V_H_H-IgG Fc fusion protein (**Table 3**; **Figure 4**). L8G12 (bin 1) and L8D9 (bin 3) also demonstrated >10-fold increase in agglutinating activity when expressed as an IgG fusion protein. In all instances, antibody-mediated spirochete agglutination correlated with a corresponding increase in *B. burgdorferi* outer membrane permeability, as reflected by elevated levels of propidium iodine (PI) uptake (**Table 3**; **Figure 4**).

**Figure 4.**
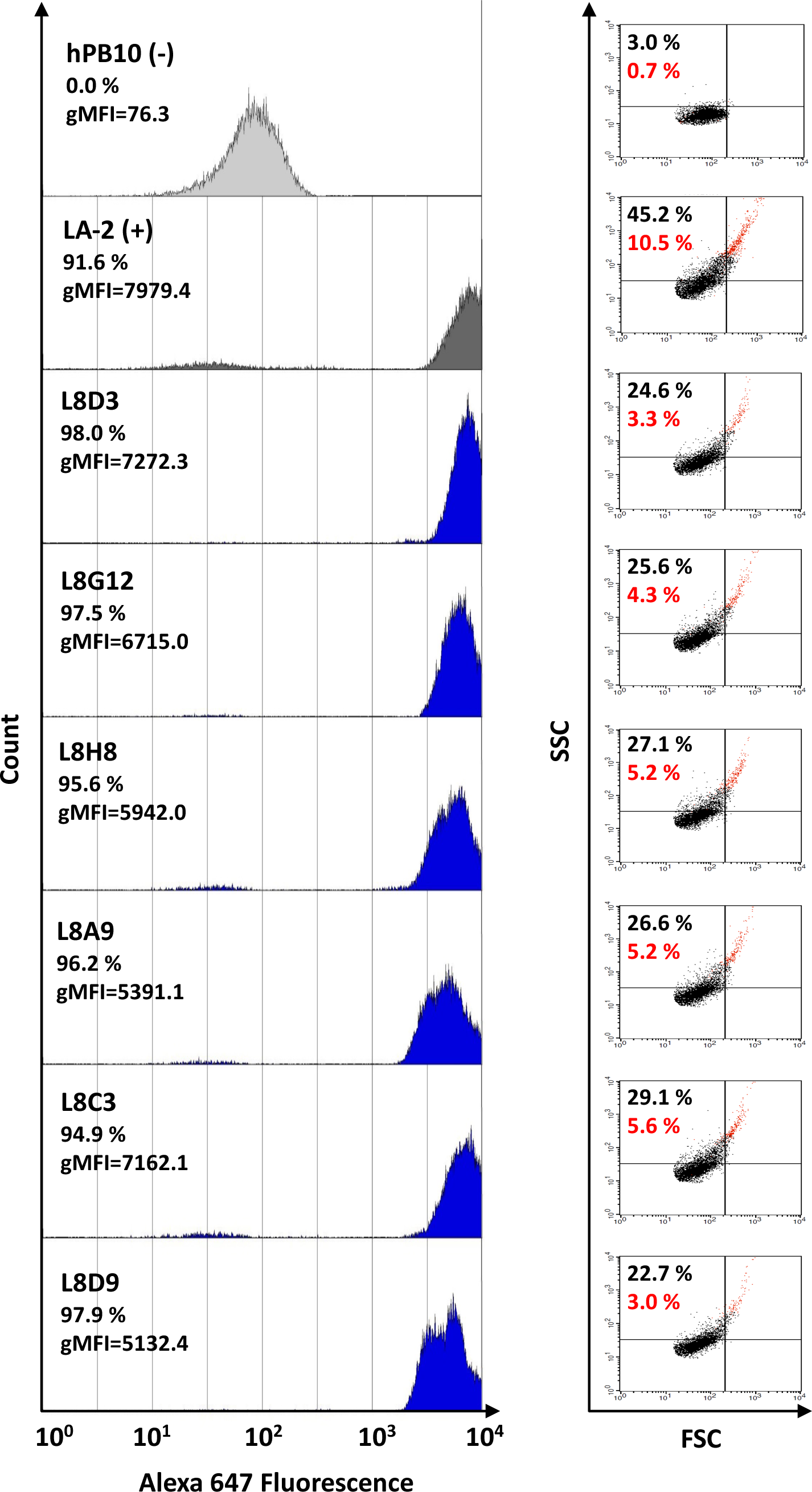
Surface labeling, agglutination and membrane permeability associated with OspA V_H_H-IgG. Flow cytometry analysis of live *B. burgdorferi* strain B31 incubated with OspA V_H_H-IgG Fcs, where an Alexa 647 fluorescent-labeled anti-human IgG secondary antibody was used to detect bound V_H_H-IgG Fcs. (**Left panel**) Representative histogram analysis of *B. burgdorferi* surface labeling by control and experimental MAbs. PB10 IgG and LA-2 IgG were used as negative and positive controls, respectively. The geometric mean fluorescence intensity (gMFI), and percent of positive events for Alexa 647 fluorescence are indicated. (**Right panel**) Corresponding forward-scatter (FSC)/side-scatter (SSC) dot plots. The percent of events that are agglutinated is indicated (black) and was calculated from the sum of events with increased FSC and SSC, in the upper-left, upper-right, and lower-right quadrants, relative to total events counted (20,000). The percent of events positive for PI staining, indicating membrane damage, is shown in red.

**Table 3.** Functional activity associated with monovalent (V_H_H) and bivalent (VHH-Ig) OspA antibodies.

| V<sub>H</sub>H | Bin | VHH | VHH-IgG1 Fc |  |  |  |  |
| --- | --- | --- | --- | --- | --- | --- | --- |
|  |  | % Agg<sup>a</sup> | % Agg<sup>a</sup> | % Binding<sup>b</sup> | MFI<sup>b</sup> | %PI<sup>+c</sup> | CDK (nM)<sup>d</sup> |
| L8D3 | 1 | 0.28 +/- 0.49 | 17.33 +/- 6.11 | 98.51 +/- 0.76 | 6894.89 +/- 425.76 | 2.23 +/- 0.47 | 2.5 |
| L8G12 | 1 | -0.03 +/- 0.05 | 18.02 +/- 6.48 | 98.23 +/- 1.05 | 6168.56 +/- 664.89 | 3.27 +/- 0.44 | 1.25 |
| L8H8 | 1 | 10.12 | 20.70 +/- 4.88 | 96.26 +/- 0.92 | 5338.44 +/- 745.55 | 4.53 +/- 0.16 | 2.5 |
| L8A9 | 3 | 7.74 +/- 0.93 | 21.99 +/- 2.28 | 97.39 +/- 1.75 | 5374.96 +/- 85.12 | 4.18 +/- 0.38 | 1.25 |
| L8C3 | 3 | 12.59 +/- 2.33 | 25.25 +/- 1.25 | 96.57 +/- 2.38 | 7141.05 +/- 78.21 | 5.52 +/- 0.98 | 1.25 |
| L8D9 | 3 | 0.65 +/- 0.50 | 16.47 +/- 4.57 | 98.46 +/- 0.80 | 4802.12 +/- 359.14 | 2.37 +/- 0.19 | 1.25 |

<sup>a</sup>, Agglutination (%) with background subtracted, n=3 (except L8H8, n=1); <sup>b</sup>, *B. burgdorferi* surface binding, n=2; <sup>c</sup>, Propidium iodide positive cells, n=2; <sup>d</sup>, Antibody-mediated, complement-dependent killing (CDK) assay are shown (EC<sub>50</sub>) n=3. +/- values are standard deviations. By comparison, the EC<sub>50</sub> values of 857-2 and LA-2 were 1.25 and ~0.75 nM, respectively.

The ability of OspA antibodies to elicit borreliacidal activity is considered a correlate of protection in Lyme disease (50). Therefore, we next assessed the OspA-specific V_H_Hs and V_H_H-IgG Fc fusion proteins for the ability to promote complement-mediated killing of a fluorescent *B. burgdorferi* B31 reporter strain (33). As expected, none of the monovalent V_H_Hs were borreliacidal (**data not shown**). However, all six V_H_H-IgG Fc fusion proteins were borreliacidal. The three bin 1 V_H_H-IgGs (L8D3, L8G12, L8H8) were tested side-by-side with 857-2, while the three bin 3 V_H_H-IgGs (L8A9, L8C3, L8D9) were compared to LA-2. The six V_H_H-IgGs tested had EC_50_ values between 1.25 and 2.5 nM (**Table 3**; **Figure 5**). By comparison, 857-2 and LA-2 had EC_50_ values of 1.25 and ∼0.75 nM, respectively (**Figure 5**). Collectively, these results demonstrate the ability of alpaca-derived V_H_Hs to recognize native OspA on the surface of live *B. burgdorferi* and promote spirochete agglutination, outer membrane damage, and complement-mediated borreliacidal activity.

**Figure 5.**
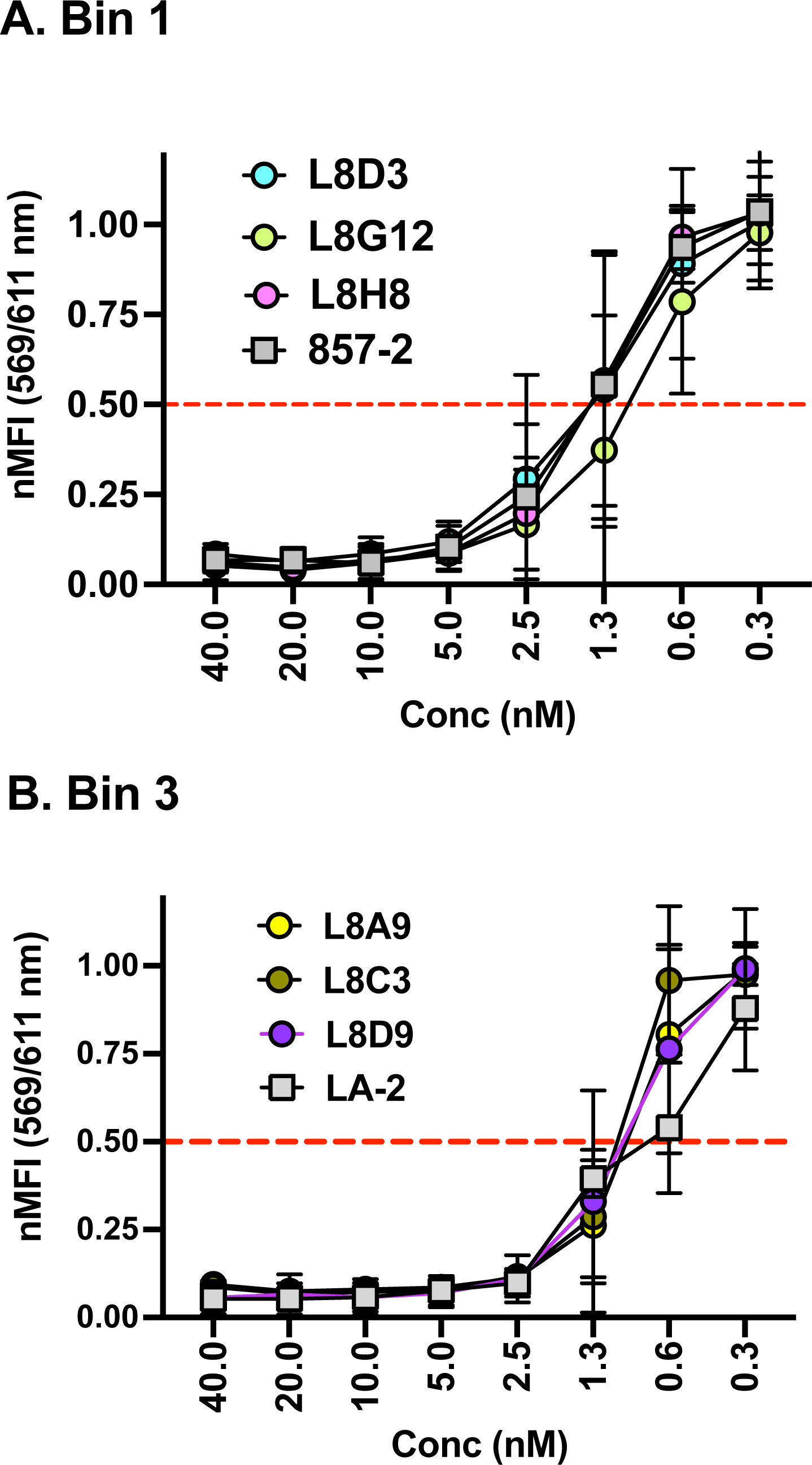
Complement-dependent borreliacidal activity imparted by V_H_H-IgG Fc. Reporter strain GGW979 was mixed with serial dilutions of each V_H_H-Fc IgG1 (A) Bin 1 and (B) Bin 2 diluted in phenol-free BSKII supplemented with gentamicin (40 μg/mL) and 20% human complement. 857-2 and LA-2 IgG1 MAbs were included in each assay to serve as positive controls for Bin 1 and Bin 3, respectively. Spirochete viability was measured 3 days later as detailed in the Materials and Methods. Data are the results of three biological replicates with SD. EC_50_ values were determined by the lowest dilution of antibody resulting in 50% reduction in MFI relative to normalized controls.

### Cross-reactivity of V_H_Hs with OspA serotypes ST1-7 reveal additional levels of epitope diversity

There are seven OspA serotypes (ST1-7) within *B. burgdorferi sensu latu* strains associated with disease (**Figure 6A**) (14, 29). The serotypes were originally defined based on reactivity profiles with a panel of OspA MAbs. LA-2 (bin 3), for example, recognizes OspA ST1 but not ST 2-7, while 857-2 (bin 1) is predicted to react with all 7 serotypes (32). To assess the specificity of the alpaca-derived V_H_Hs, the three bin 1 V_H_H-IgGs (L8D3, L8G12, L8H8) and three bin 3 V_H_H-IgGs (L8A9, L8C3, L8D9) were evaluated in an OspA ST 1-7 indirect ELISA (**Figure 6B**). As predicted, 857-2 reacted with all seven OspA serotypes, while LA-2 reacted with only ST-1. The V_H_Hs displayed unique reactivity patterns relative to 857-2 and LA-2. Within bin 1, for example, L8D3 recognized ST1 exclusively, while L8G12 and L8H8 recognized ST 1, 2, 4 and 5. Within bin 3, L8D9 reacted exclusively with serotype 1 (like LA-2), but L8A9 and L8C3 recognized ST1 plus serotype 6 (L8A9) or serotype 4 (L8C3). These results reveal a greater degree of B cell epitope diversity on OspA than previously recognized and have important implications for multivalent vaccine design.

**Figure 6.**
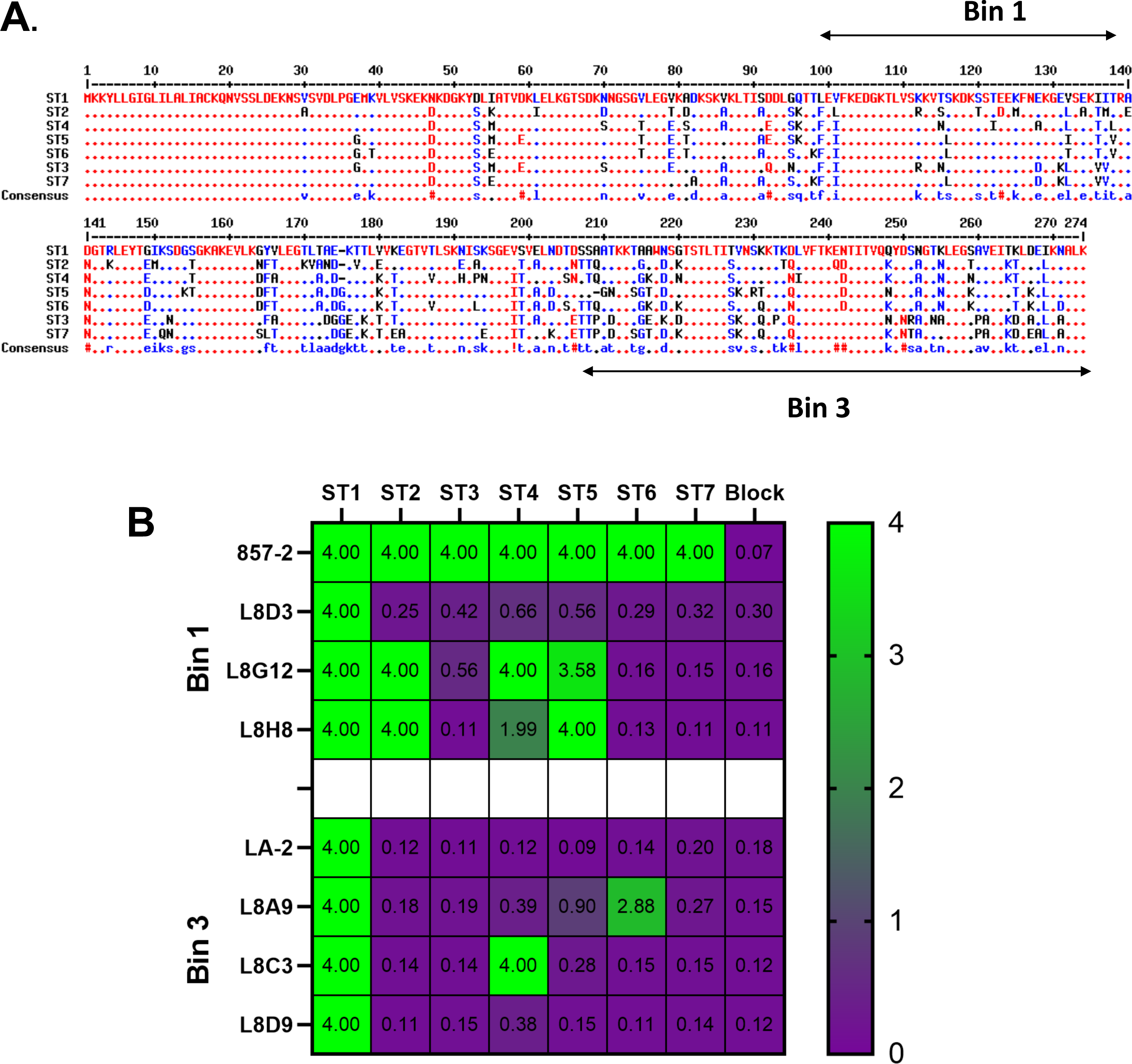
V_H_H-IgG Fc recognition of OspA serotypes by ELISA. (A) Amino acid alignment of the seven OspA serotypes (ST1-7). Dots denote amino acid identity with ST1; one letter amino acid codes shown in locations that differ from ST1. (**B**) Microtiter plates were coated with indicated OspA serotypes (horizontal labels) or block solution only (control), then probed with indicated Bin 1 (857-2, L8D3, L8G12, L8H8) or Bin 3 (LA-2, L8A9, L8C3, L8D9) antibodies (vertical labels) and developed as described in the Materials and Methods. The heat map scale (purple to green) is shown on the right and the numbers refer to absorbance values upon addition of stop solution. Deep purple (0) represents no binding, while lighter green (4) represents strong binding.

## DISCUSSION

Single-domain antibodies (V_H_Hs) have emerged as invaluable tools and reagents in the development of next-generation diagnostics, therapeutics, and vaccines for a range of infectious diseases (40). With this in mind, we sought to generate a diverse collection of V_H_Hs against the Lyme disease vaccine antigen, OspA, as a means to better define the conserved and variable epitopes on the different OspA serotypes associated with borreliacidal activity. We constructed an immune V_H_H-phage display library from two OspA immunized alpacas and subjected the library to rounds of affinity enrichment on recombinant OspA serotype 1 (ST1). The screen yielded 21 unique OspA-specific V_H_Hs with a range of binding affinities, epitope specificities, serotype reactivities, and functional activities *in vitro*. A subset of V_H_Hs were expressed as human IgG1-Fc fusion proteins and shown to promote complement-independent *B. burgdorferi* agglutination, as well as complement-dependent borreliacidal activity. In addition to expanding our understanding of functional B cell epitopes on OspA, we expect that this unique collection of V_H_Hs will have applications for pre-clinical and clinical Lyme disease vaccine development.

The 21 V_H_Hs described in this study have epitope reactivity profiles that are both similar and different from previously reported OspA-specific mouse and human MAbs (17, 19, 26–29, 33, 34, 43, 51). On the one hand, the V_H_Hs fell within two previously described epitope bins based on competition with human MAb 857-2 (bin 1) and mouse MAb LA-2 (bin 3). On the other hand, the V_H_Hs displayed unique reactivity profiles within those respective bins. For example, within bin 1, 857-2 recognized all seven OspA serotypes, while L8G12 and L8H8 were reactive with serotypes 1, 2, 4 and 5; L8D3 was only reactive with serotype 1. This result is consistent with pioneering work by Wilske and colleagues demonstrating that there are conserved, partially conserved, and serotype-specific epitopes within OspA’s central β-sheet (strand 8-14) (29).

Competition assays with three different bin 3 MAbs (LA-2, 319-44, 3-24) enabled us to localize epitopes recognized by seven different V_H_Hs to OspA β-strands 16-18 (residues 195-227). However, even within this relatively restricted region of OspA, there was notable epitope diversity, as exemplified by differential serotype reactivities of L8A9 and L8C3 as compared to LA-2. Specifically, LA-2 recognized serotype 1 exclusively, while L8A9 and L8C3 were reactive with serotypes 1, 4, and 6. This observation challenges the notion that the C-terminal region of OspA elicits only serotype-specific antibody responses (28, 52).

From the standpoint of V_H_H clonal diversity, we identified five distinct antibody families that collectively accounted for more than half the 21 V_H_Hs. The L8H8 family is notable for having the most members (four) recovered in our current screening protocol, as well as the V_H_H with the highest affinity for OspA (L8H8; 0.28 nM). As expected, the three L8H8 family members we subjected to epitope mapping by HX-MS had virtually identical protection profiles. However, within the family, the dissociation constants ranged by at least an order of magnitude from 0.28 nM (L8H8) to >2 nM (L8E1), likely due to variations in CDR1 and CDR2 interactions with OspA. This novel collection of antibodies with identical epitope specificities but varying binding affinities can be used as a toolkit to investigate the open question of the relative contribution of binding affinity in OspA antibody effector function. As a case in point, studies are planned to evaluate the L8H8 family of V_H_Hs as well as V_H_H-IgGs for the ability to block transmission of *B. burgdorferi* in a mouse model of tick-mediated infection, with the expectation that there will be a direct relationship between binding affinity and protection. Indeed, as shown in Table 1, the contribution of binding affinity on the ability of the L8H8 family of V_H_Hs to promote spirochete agglutination is already somewhat evident.

While the V_H_Hs and V_H_H-IgGs described in this study have yet to be tested for *B. burgdorferi* transmission blocking activity in a mouse model, we would predict, based on *in vitro* activities, that at least a subset will be effective *in vivo*. Most significant in our minds was the observation that a subset of monovalent V_H_Hs and all six bivalent V_H_H-IgGs could promote spirochete agglutination and induce alterations in outer membrane permeability. We have argued that such activities, should they occur in the context of the tick midgut, would impair the ability of spirochetes to migrate across the midgut epithelium and onto the salivary glands (46). The V_H_H-IgGs (but not the monovalent V_H_Hs themselves) were also extremely effective at promoting complement-mediated killing of B*. burgdorferi*. While the role of complement-mediated killing in transmission inhibition within the tick remains an open question, OspA antibody-mediated complement-dependent killing *in vitro* does correlate with protection *in vivo* (50). Thus, we predict that the six V_H_H-IgGs described in Table 3 will likely prove effective *in vivo*.

We envision at least two potential applications of the V_H_Hs to Lyme disease vaccine development. First is their use in “equivalency” assays. For example, in the pivotal clinical trial associated with the LYMErix vaccine, a competitive ELISA with LA-2 was used as a surrogate measure of protective antibody titers limited to OspA serotype 1 (24). Similar, albeit more sophisticated, competition assays using a panel of V_H_Hs directed against known protective epitopes on OspA serotypes 1-7 could be employed for the evaluation of future multivalent Lyme disease vaccines. Second, the OspA V_H_Hs could be employed in identity testing, epitope integrity analysis, and release assays associated with vaccine manufacturing and mandated by regulatory agencies. Identity testing will be especially relevant when evaluating recombinant or nucleic acid-based hexavalent and even heptavalent Lyme disease vaccines (6). We have also described other antibody-based applications for vaccine development that may be pertinent to Lyme disease (53).

## Materials and Methods

### Alpaca immunization and V_H_H phage display library preparation

Two alpacas (*Vicugna pacos*) were immunized four times, each about three weeks apart, with various combinations of recombinant OspA serotype 1 from *B. burgdorferi* strain B31 [UniProt P0CL66] and the veterinary vaccine RECOMBITEK^®^ Lyme (Boehringer Ingelheim). The first immunization consisted of a full dose of RECOMBITEK^®^ only. The next vaccination the alpacas received a half dose of RECOMBITEK^®^ and 200 µg of rOspA. The final two vaccinations consisted of a half dose of RECOMBITEK® and 100 µg of rOspA. Following the final immunization, both animals had OspA antibody endpoint titers > 500,000. Five days following the final immunization, lymphocytes were isolated from whole blood and used to generate a V_H_H M13 phage-display library, as described (54, 55). The library passed all QC testing and was estimated to contain ∼1.3 x 10^7^ independent clones.

### Recombinant OspA

Recombinant OspA (non-lipidated) ST1 from B31 [NCBI reference WP_010890378.1] was expressed in *E.coli* and purified as described (33, 34). Purified, recombinant OspA serotypes 2-7 were kindly provided by Dr. Meredith Finn (Moderna, Inc).

### V_H_H Identification and Expression

The V_H_H M13 phage-display library was subjected to two rounds of affinity enrichment (panning) against immobilized recombinant OspA on Nunc-Immunotubes™ (Thermo Fisher Scientific, Waltham, MA) using protocols described elsewhere (56). The first panning was low stringency (10 ug/mL coat), after which phages were eluted with glycine HCl [pH 2.2] and then amplified in *E. coli*. The phages were then subjected to a second round of screening at high stringency (1 ug/mL coat). After the second round of panning, 95 clones were chosen at random and grown overnight at 37° C in a 96 well plate. Replica cultures in 96 well plates were grown to log phase, then induced overnight with IPTG (3 mM). Resulting supernatants were assayed by ELISA for reactivity with rOspA. 74 clones were found to bind to OspA, and all were then DNA sequenced. 21 V_H_Hs, belonging to 12 different sequence families, were found to be sufficiently unique to characterize further. The DNA coding region for these V_H_Hs was restriction digested out of the phagemid vector and inserted into a pET-32b vector for expression as a recombinant thioredoxin fusion protein containing a His-Tag in the linker region, and an E-tag for detection at the C-terminus. V_H_Hs were transformed into and expressed in Rosetta Gami 2 (DE3) pLacI competent *E. coli* (Millipore Sigma), induced with IPTG (1 mM), and purified in a nickel column. Concentration was determined by OD_280_ and the recombinant protein extinction coefficient.

### ELISA

OspA was coated overnight in 96-well immunoplates at 1 ug/mL in 100 uL of PBS. The following morning, the wells were blocked for 2 hours with 2% goat serum in PBS with 0.1% Tween 20 (PBST). During blocking, V_H_Hs were 5-fold diluted in PBS in a separate 96-well non-binding plate, starting at 10 uM. The V_H_Hs were then applied to the OspA coated plate for 1 hour and allowed to bind. After washing the wells with PBST, HRP-conjugated anti-E-tag secondary antibody (Bethyl Labs, Waltham MA) was added to the well for 1 hour to detect bound V_H_Hs. After a final wash with PBST, 100 uL SureBlue TMB (SeraCare) was added to the wells for about 10 minutes to visualize binding. The colorometric reaction was quenched with 1M phosphoric acid, and absorbance at 450 nm was measured on a SpectraMax iD3 plate reader (Molecular Biosystems) using SoftMax Pro version 7.1 software.

### Biolayer Interferometry (BLI)

BLI experiments were carried out using an Octet RED96e Biolayer Interferometer (Sartorius AG, Gottingen, Germany) using the Data Acquisition 12.0 software. Raw sensor data was loaded into the Data Analysis HT 12.0 software for analysis. Biotinylated OspA (5 µg/mL) in PBS containing 2% w/v BSA was captured onto streptavidin biosensors (#18-5019, Sartorius) for 5 min. After 5 minutes of baseline in buffer, sensors were then exposed to a 2-fold dilution series of V_H_H, ranging from 200 to 3.125 nM, for 5 minutes to allow association. The sensors were then immediately dipped into wells containing buffer alone for 30 minutes to allow dissociation of the V_H_H. An eighth sensor was also loaded with Biotinylated OspA, but was not exposed to V_H_Hs, and was thus used as a background drift control, and subtracted from the other sensor data. After each V_H_H, the OspA-coated sensors were completely regenerated by a 30 second cycle consisting of three repeats of 5 sec in 0.2 M glycine (pH 2.2) and 5 sec in buffer. Sensor data was fit to a 1:1 binding model.

For competition experiments biotinylated OspA (5 µg/mL) in buffer was captured onto streptavidin biosensors for 5 minutes. After 3 minutes of baseline in buffer, sensors were then exposed to a primary mAb at a concentration of 15 µg/mL in buffer for 10 minutes to permit the association signal to saturate. The sensors were then immediately dipped into wells containing competitor V_H_H (1 uM) for 10 min. After each primary-secondary pairing, the sensors were regenerated by a 30 second cycle consisting of three repeats of 5 sec in 0.2 M glycine (pH 2.2) and 5 sec in buffer. The total binding signal (in nm) obtained for the secondary V_H_H, from the end of the primary mAb, was then recorded for each specific primary-secondary pair. The data for each V_H_H was normalized to the binding signal for that V_H_H vs mAb 212-55, which no V_H_H competed with, then plotted as a heat map using GraphPad Prism 9.

### HX-MS

Differential hydrogen exchange-mass spectrometry (HX-MS) was used to identify regions of OspA exhibiting altered amide hydrogen exchange kinetics in the presence of an excess of each V_H_H using methods fully described previously (33, 44). In brief, OspA alone or in the presence of molar excess V_H_H was diluted ten-fold with 20 mM phosphate, 100 mM NaCl, pH 7.40 buffer containing deuterium oxide. After various intervals of exchange ranging between 20 s and 24 hr, the exchange reaction was rapidly quenched by acidification. HX-MS measurements were completed either in triplicate at five exchange times (referred to as “complete”) or single measurements at three exchange times (referred to as “screening”). The quenched samples were rapidly digested with pepsin to yield deuterium-labeled OspA peptides. The deuterium incorporation was measured by LC-MS. Differences in hydrogen exchange were quantified as the mean exchange by bound OspA minus mean exchange by free OspA. Since peptides of different lengths contain different numbers of amide hydrogens, the results were normalized based on the amount of deuteration in maximally deuterated control samples. The resulting quantity, Δ*HX̅* represents this mean fractional difference; negative values indicate slower hydrogen exchange by bound OspA.

### Surface binding, membrane integrity, and agglutination analysis of *B. burgdorferi*

*B. burgdorferi* strain B31 (ATCC) was cultured in BSK-II media (minus gelatin) at 33°C with 2.5% CO_2_ (57). Cultures at mid-log phase were diluted 1/10 in medium and grown at 23°C to early-log phase to induce high levels of OspA expression (46). Bacteria were collected by centrifugation (3,300 x *g*), washed with PBS, resuspended in BSK II medium (minus phenol red), and allowed to recover at room temperature for 30 min. A total of 5x10^6^ cells in 50 µl were incubated with an OspA-specific VHH or VHH-IgG Fc at a final concentration of 10 µg/ml at 37 °C for 1 h. Incubation with 10 µg/ml of an unrelated ricin-specific VHH (V8B3) or IgG (PB10) were included as negative controls, while chimeric LA-2 IgG1 was run as a positive control (33). Reaction volumes were increased with the addition of 450 µl of BSK II medium (minus phenol red) and incubated at 37°C for 30 min with either a 1/500 dilution of PE-labeled anti-6xHis tag mouse MAb (Biolegend, San Diego, CA) to detect VHHs or Alexa Fluor 647-labeled goat anti-human IgG (H+L) (Invitrogen) to detect VHH-IgG Fcs. All mixtures were transferred into a 5 mL round bottom polystyrene test tube (Corning) with 250 µl of PBS. For VHH-IgG Fc reactions, 0.75 µM propidium iodide (PI) (Sigma-Aldrich) was added immediately before analysis to measure membrane integrity. All samples were analyzed on a BD FACSCalibur flow cytometer (BD Biosciences). Bacteria were gated on forward scatter (FSC) and side scatter (SSC) to assess aggregate size and granularity, and 20,000 events were counted. Alexa Fluor 647 labeling (FL4), PE labeling (FL3) or PI staining (FL3), and agglutination (FSC/SSC) were measured using CellQuest Pro (BD Biosciences). Agglutination was calculated as the sum of events in the upper-left, upper-right, and lower-right quadrants relative to total events (46).

### Borreliacidal assays

V_H_H-IgG Fcs were assessed for complement-dependent borreliacidal activity via fluorescence-based serum bactericidal assay essentially as described (33). Slight modifications to the previous assay include the mCherry open reading frame (ORF) within the reporter plasmid (pGW163) was replaced with a codon-optimized variant of mScarlet-I engineered for efficient expression in *Borrelia* species (G. Willsey, N. Mantis, *manuscript in preparation*). The resulting plasmid, pGW189, was subsequently transformed into *B. burgdorferi* B31-5A4 following established protocols (58). The resulting IPTG-inducible mScarlet-I viability reporter strain was designated GGW979.

For each assay, glycerol stocks of GGW979 were thawed at RT and transferred to sterile 50 ml centrifuge tubes containing 45 ml of BSK-II medium supplemented with 40 µg/ml of gentamicin. The cultures were then incubated at 33°C without agitation for three days. On the day of the assay, spirochetes were collected via centrifugation, the culture media was removed, and the cells were then resuspended at 3 x10^7^ spirochetes per ml in phenol-free BSKII medium supplemented with gentamicin (40 μg/mL) and 20 % human complement sera (SigmaAldrich). The spirochetes were then mixed 1:1 v/v with serial dilutions of each V_H_H-Fc IgG1 antibody that had been prepared in phenol-free BSKII supplemented with gentamicin (40 μg/mL) and 20 % human complement. Following sample addition, each reaction contained and 5x10^6^ spirochetes per well and 40 nM and 0.325 nM of each antibody. 857-2 and LA-2 IgG1 MAbs were included in each assay to serve as positive controls (33). Untreated/non-induced and untreated/IPTG-induced controls were similarly included to determine baseline and peak fluorescence.

Following sample addition, assay plates were incubated overnight at 37 °C with 5 % CO2. After 18-20 h, 1 mM IPTG was added to each well (minus the untreated control) to induce expression of the fluorescent reporter in surviving spirochetes. Assay plates were then returned to the incubator. Forty-eight hours later, Median Fluorescence Intensity (MFI) was recorded three times per plate at 569 nm (Ex)/611 nm (Em) using a SpectraMax iD3 microplate reader (Molecular Biosystems) and SoftMax Pro version 7.1 software. The resulting values were averaged, and the data was then normalized using the untreated (-IPTG) and untreated (+IPTG) controls to set baseline (0) and peak (100) MFI. Following normalization, the data was analyzed using GraphPad Prism Version 9.0. Data reported encompasses three separate experiments, with EC_50_ values determined by the lowest dilution of antibody resulting in 50% reduction in MFI relative to normalized controls.

## Author Contributions

DJV screened the V_H_H phage display library, sequenced the V_H_Hs, conducted Octet and ST1-7 ELISA experiments, and wrote and edited the manuscript; SB cloned and expressed VHHs in *E.coli*, and performed OspA ELISA experiments; CLP conducted *B. burgdorferi* flow cytometry experiments; GGW conducted *B. burgdorferi* borreliacidal assays; JMT and CBS constructed the V_H_H phage display library; HMEH and DDW conducted HX-MS and data analysis; MJR expressed and purified all recombinant proteins; LC expressed and purified the VHH-Fc antibodies; NJM was responsible for project leadership, funding acquisition, and writing the manuscript.

## ACKNOWLEDGEMENTS

The authors gratefully acknowledge Elizabeth Cavosie (Wadsworth Center) for administrative assistance and grants management. We thank the Wadsworth Center’s Immunology Core for flow cytometry assistance and the Cell culture and media core for BSK II medium. We extend our special thanks to Drs. Meredith Finn and Chris Dold (Moderna, Inc, Cambridge, MA) for providing recombinant OspA serotypes 1-7. This work was supported by the National Institute of Allergy and Infectious Disease (NIAID), National Institutes of Health, Department of Health and Human Services, Contract No. 75N93019C00040 (PI/PD Mantis).

